# A structure-based approach towards identification of inhibitory fragments for eleven-nineteen-leukemia protein (ENL) YEATS domain

**DOI:** 10.1101/466284

**Authors:** David Heidenreich, Moses Moustakim, Jurema Schmidt, Daniel Merk, Paul E. Brennan, Oleg Fedorov, Apirat Chaikuad, Stefan Knapp

## Abstract

Lysine acetylation is an epigenetic mark that is principally recognized by bromodomains and recently structurally diverse YEATS domains also emerged as readers of lysine acetyl/acylations. Here we present a crystallography-based strategy and the discovery of fragments binding to the ENL YEATS domain, a potential drug target. Crystal structures combined with synthetic efforts led to the identification of a sub-micromolar binder, providing first starting points for the development of chemical probes for this reader domain family.

---

Post-translation modification at lysines is one of the key mechanisms that regulates epigenetic signaling. Lysine acetylation (Kac) is one of the most common epigenetic “marks”, which is specifically recognized by bromodomains and some double PHD finger (DPF) domains^1-2^. Recently, YEATS domains emerged as a third class of histone acetylation readers which are present in four human proteins: ENL (MLLT1), YEATS2, AF9 (MLLT3) and glioma amplified sequence 41 (GAS41 or YEATS4). Interaction with acetylated histones was first demonstrated for AF9^3^ and subsequently also confirmed for the remaining family members^4-6^. Interestingly, YEATS domains display an expanded reader activity, also recognizing other types of lysine modifications including propionylation, butyrylation and crotonylation^3, 7-9^.

The YEATS domain constitutes ^∼^120-140 amino acids and is evolutionarily conserved from yeast to human^10^. It exhibits an immunoglobulin-like topology with an elongated β-sheet sandwich core capped by one or two short helices^3, 11^. The binding pocket is constructed by three loops emanating from the Ig fold. A number of conserved aromatic residues shape a flat, extended architecture of the binding groove, that is capable of recognizing acyl-lysine containing sequences^3-7, 9^. The proteins harboring the YEATS reader domain are often associated with histone acetyl transferase (HAT) and chromatin-remodeling complexes^12-13^, implicating their diverse roles in the regulation of chromatin structure, histone acetylation, gene transcription, stress signaling, mitotic progression and DNA damage response^3, 13-15^. In addition, the ability to preferentially recognize other lysine acylation marks suggests that the YEATS family proteins might exert differential regulatory functions than the prototypical Kac reader families with their own cognate targets^16^.

Dysfunction of YEATS proteins has been linked to diseases, notably cancer. For instance, the fusion of AF9 or ENL and human mixed lineage leukemia (MLL) proteins are frequently found in acute myeloid leukemia, and these fusions constitute oncogenes that are drivers of this highly aggressive cancer^5, 17-18^. In addition, GAS41, a common subunit of SRCAP (Snf2 related CREBBP activator protein) and Tip60 HAT complexes, is a growth-promoting protein^12, 19^. These roles suggest that YEATS proteins are potential targets for drug development. Indeed, two recent studies identified the ENL (MLLT1) YEATS domain as a compelling target in AML^5^, ^17^.

Development of chemical probes targeting the acetyl-lysine readers, the bromodomain family, provided significant insight into the biological function of these proteins and their potential as drug targets^20-22^. However, in contrast to bromodomains, no inhibitors of YEATS have been reported to date. This prompted us to identify chemical scaffolds that can interact with the binding pocket of this acetyl-lysine reader family, focusing on oncogenic ENL. In a similar manner to bromodomains, we chose a structure-based approach to identify initial fragment binders^23-26^. We predicted an essential chemical moiety that could mimic β-sheet type hydrogen bonding pattern of acylated lysine, and established a small fragment-like library. Using structure-based approaches as well as biophysical characterization such as thermal shift and isothermal calorimetry (ITC), we identified potential inhibitory ligands for ENL, which may provide chemical starting points for further development of potent inhibitors for this protein as well as the other members of the histone acylation reader YEATS family.

To date all available crystal structures of ENL and other YEATS proteins have been solved in complexes with peptides. We therefore determined the apo-structure of ENL to expand our knowledge on a non-liganded form of the protein. In the ENL apo-structure, all structural elements were well defined by electron density, including loop 1 (L1), 4 (L4) and 6 (L6) that defined the recognition site for acylated lysine (Figure 1A and Supplementary Figure S1). This suggested that the binding pocket is well-structured prior to the binding of the ligands. Nonetheless, surprisingly, the side chain of Y78 located on L6, which is together with F28 and F59 part of the conserved aromatic acyl-lysine binding triad, exhibited two orientations not observed previously in the peptide-complexed structures. The ‘in’ conformation with the side chain positioned on top of the binding site resembled the orientation observed in ENL-K_ac_27H3 complex, while the ‘out’ conformation exhibited a 90° side chain rotation. This conformation was stabilized by a hydrogen bond to the backbone of L1 E26. The unexpected flexibility of this tyrosine suggested therefore an intrinsic flexible nature of the binding site, interchangeable between the confined ‘in’ conformation and a more open surface when adopting the ‘out’ conformation.

**FIGURE 1.**
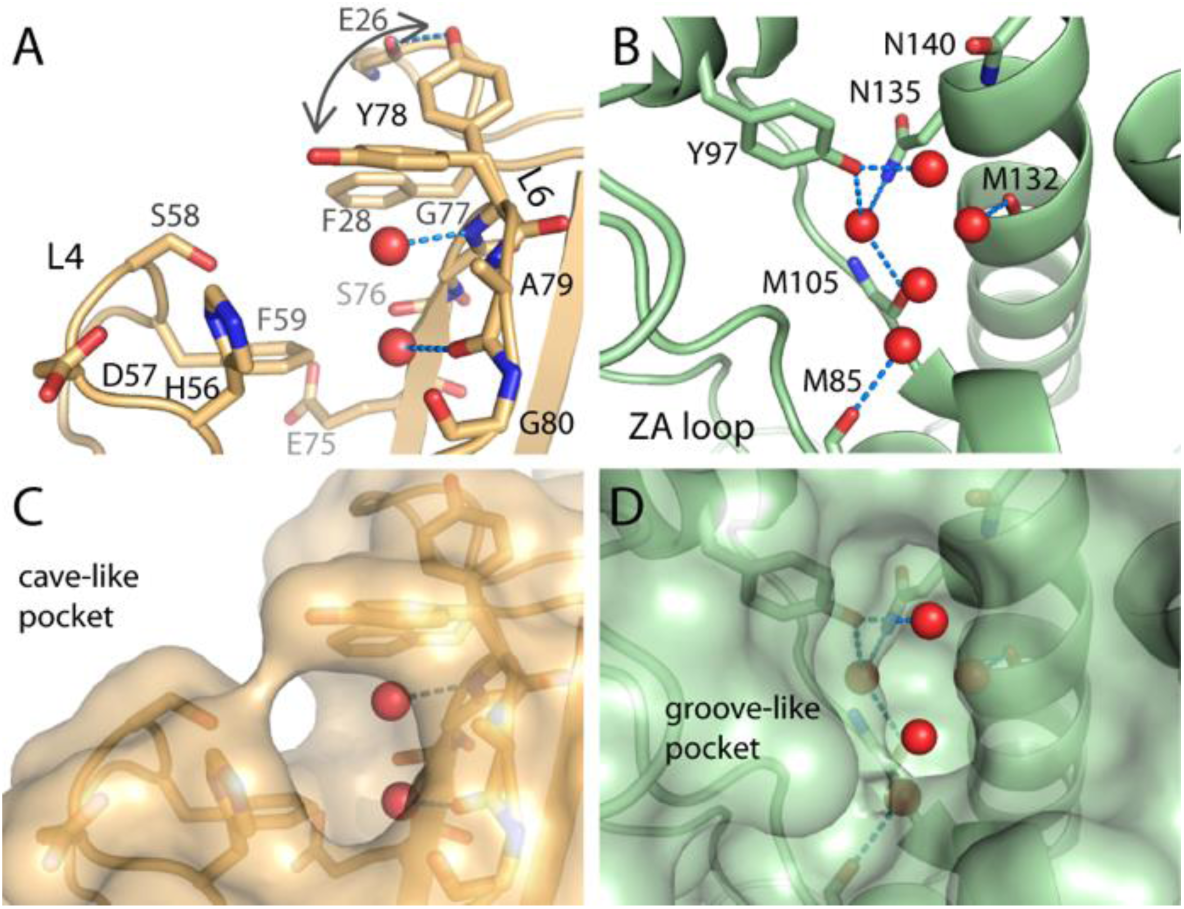
Structure of apo-ENL (pdb code, 6hq0). A) close-up details of the binding site of ligand-free ENL with Y78, a residue part of the key interacting aromatic triad, adopting two flexible conformations. Bound water molecules within the pocket are shown in red spheres. B) acetyl-lysine binding pocket of BRD4 (pdb id: 2OSS) with conserved asparagine (N140), tyrosine residue (Y97) and water network (red spheres) displayed. Surface representation of the binding sites of ENL (C) and BRD4 (D) reveals distinct pocket shapes between these two acetyl-lysine binding reader families.

Despite sharing acetyl-lysine recognition functions, comparative analyses revealed diverse features between canonical acetyl-lysine binding sites of ENL and bromodomains such as BRD4^2^ (Figure 1A and 1B). First, the two families harbored distinct primary, secondary and tertiary structure and therefore have diverse acetyl-lysine binding sites. In bromodomains, the acetyl moiety is anchored by hydrogen bonds to a conserved asparagine and via a water to a tyrosine residue (N140 and Y97 in BRD4)^2, 27^. In contrast, in ENL the backbone of loop 6 and the aromatic triad (F28, F59 and Y97 in ENL) mediate this recognition process^3, 5^. In addition, the two readers exploited different structural elements towards a formation of the pocket. Two helices and the ZA loop, acting as a selectivity filter for both protein partners and small molecule inhibitors, contributed to a deep, groove-like construction in bromodomains^2^, which was in contrast to a surface-exposed, tunnel-like pocket formed through three loop elements and the side chains of the aromatic triad in ENL (Figure 1C and 1D). These differences resulted also in diverse patterns of conserved water molecules present in the pockets. Typically five water molecules conducting a complex hydrogen bond network in bromodomains^28-29^ compared to two structural waters present in ENL located adjacent to Y78 and A79 backbones (Figure 1).

Although the binding site was pre-existent in the apo form, the unexpected movement of Y78 raised a possibility of some degrees of the flexibility of ENL. We next determined the K_ac_-complexed structure, in which the electron density conceivably allowed a placement of the weak affinity acetyl-lysine ligand (Figure 2A). The acetyl moiety displaced one water molecule for the formation of the only hydrogen bond between its carbonyl group with Y78 amine backbone. Superimposition of this structure with the ENL-peptide complex^5^ revealed highly conserved binding modes of acetyl-lysine and peptides. Nonetheless, detailed analyses demonstrated otherwise slight alterations of some residues lining the pocket in particular the reorientations of the H56 and loop4 D57 side chains (Figure 2B). This concerted movement of these two residues may modulated interactions with the adjacent S58 hydroxyl side chain and suggest an induced fit model of the acetyl-lysine interaction. However, unlike the peptide complex the presence of acetyl-lysine ligand alone was insufficient to lock the ‘in’ orientation of Y78 for the closure of the pocket, explaining low affinity of the acetyl-lysine ligand compared to the peptide, evident also by partial electron density observed for the ligand.

**FIGURE 2.**
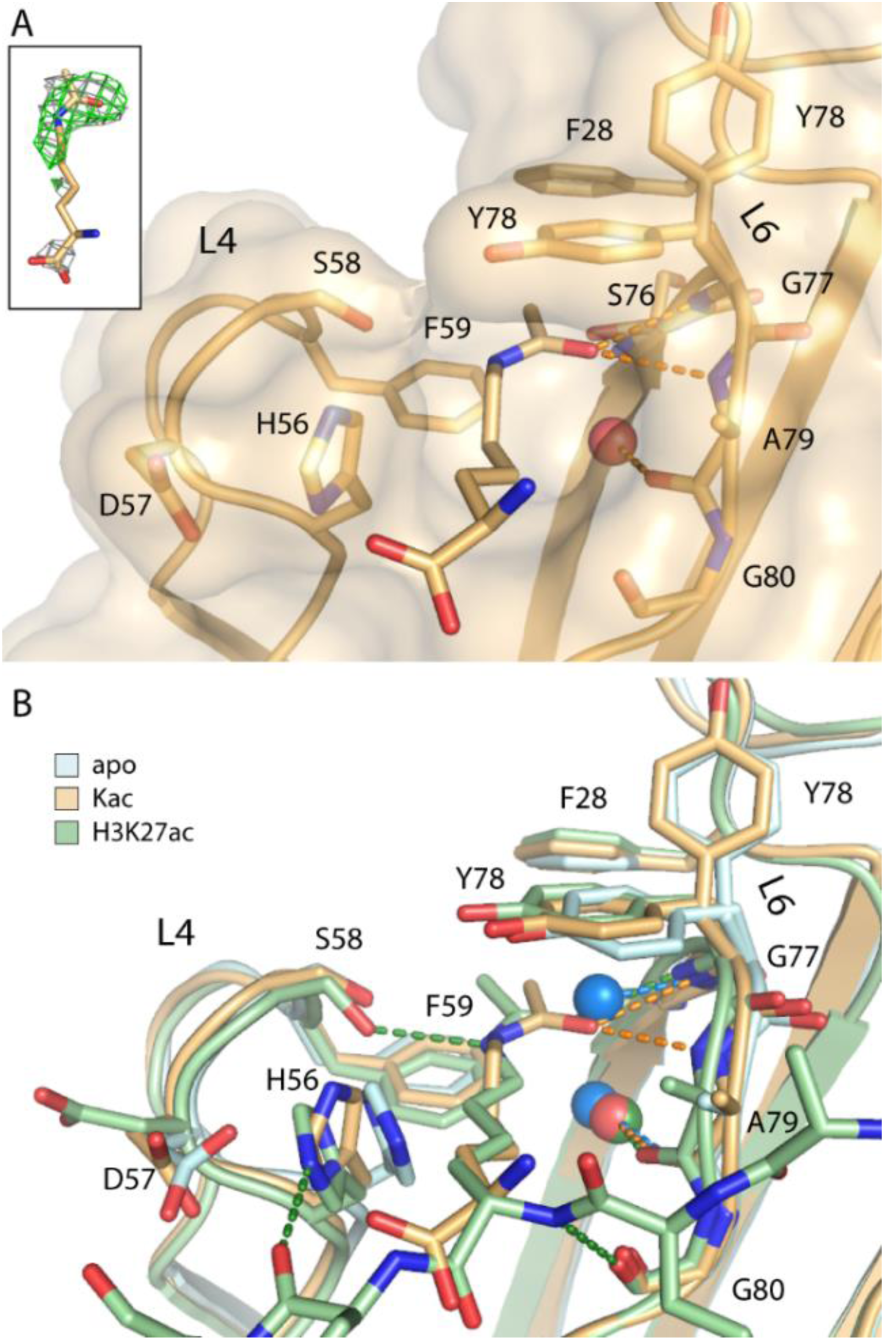
Structure of ENL in complex with acetyl-lysine. A) Detailed interaction between the bound Kac within ENL (pdb code, 6hpz). Inset shows |*F*_O_| - |*F*_C_| omitted electron density map contoured at 3σ (green) and |2*F*_O_|-|*F*_C_| refined map contoured at 1σ (grey) for the bound ligand. B) Superimposition of apo, K_ac_-bound, and H3K27ac-complexed (pdb id: 5J9S) ENL structures revealed a conserved binding mode of Kac that displaced a water for a hydrogen bond interacting with the Y78 backbone amine. The presence of the ligand or peptide induced structural alterations at Y78, H56 and D57.

Comprehensive structural analyses revealed that the distinct cave-like characteristic of the pocket in ENL, as well as in other YEATS members, may well be evolved for its compatibility with various lysine acylation. The front open adjacent to H56 and loop6 A79 provides a more restrained entrance for lysine backbone, while the rear open end cradled by F28 and F59 adopts a flat, hydrophobic environment for accepting the elongated acylation modification such as K_cr_^7, 9^ (Figure 3A). The profound structural differences between the pockets of bromodomains and YEATS domains therefore suggested that current acetyl-lysine mimetic bromodomain ligands are unlikely to bind to YEATS domains. With a largely hydrophobic and aromatic surface area of ^∼^93 Å^2^ and a pocket volume of ^∼^34 Å^3^, the ENL binding pocket is smaller than most bromodomain binding sites, yet could be sufficiently large for the development of potent inhibitors.

**FIGURE 3.**
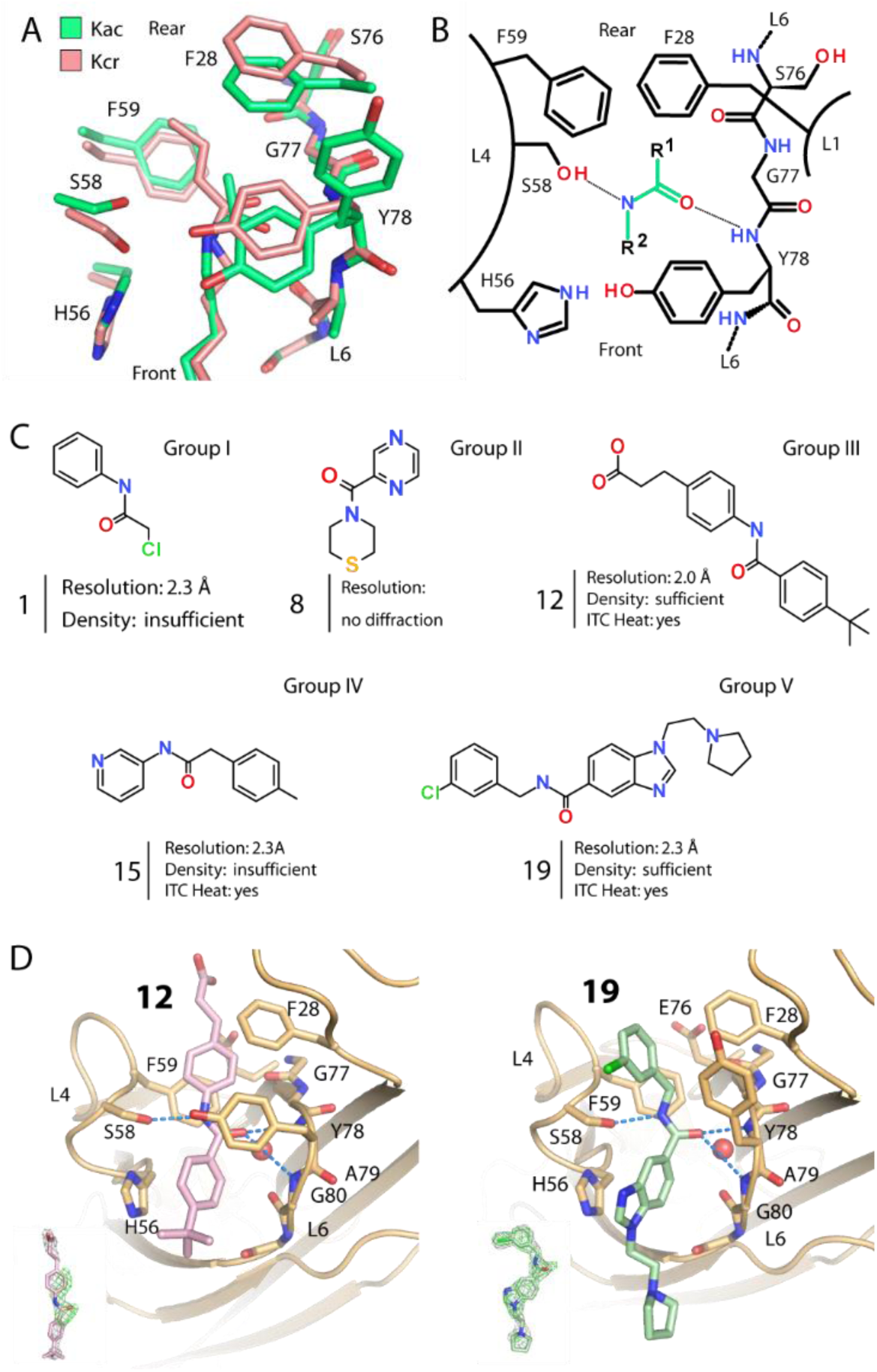
Crystallography-based screening approach. A) Analyses of the binding pockets of ENL and YEATS family reveals an open binding site. B) Schematic illustration of key residues within the pocket and the design of acetyl-lysine mimetics containing an amide core mimicking a β-sheet hydrogen bond interaction and an aromatic decoration at R^1^ or R^2^ for π-π interactions with F28 and F59. C) Examples of selected ligands. D) Structures 0f ENL in complex with **12** (top, pdb code, 6hpy) and **19** (bottom, pdb code, 6hpx). Insets show |*F*_O_| - |*F*_C_| omitted and |2*F*_O_| - |*F*_C_| electron density map contoured at 3σ (green) and 1σ (grey), respectively. Hydrogen bonds are shown as dashed lines and water molecules as red spheres.

The unique features of the ENL pocket prompted us to presume that the narrow binding site with two open ends might be able to accommodate linear molecules harboring a central carbonyl group, which were considered as initial fragment-like scaffolds (Figure 3A). For fragment selection, we postulated that suitable ligands should contain i) a central amide functional group to mimic acetyl-lysine and ii) at least an aromatic property at carbonyl (R^1^) end or, if flipped amide, nitrogen (R^2^) end to mimic μ-stacking with F28 and F59, an interaction observed for the crotonyl group (K_cr_) (Figure 3B).

Based on this assumption, we first selected a small set of 19 fragment-like compounds, which were classified into four groups based on the positions of their aromatic rings; i) only on R^2^ (**1**-7), ii) only on R^1^ (**8**), iii) on both R^1^ and R^2^ (**9-14**), iv) on R^2^ with a spacing atom prior to attachment on R^1^ (**15-18**), and v) on R^1^ with a spacing atom prior to attachment on R^2^ (**19**) (Figure 3C and Supplementary Figure S2). These compounds were initially tested using thermal shift assays, unfortunately no detectable shifts in melting temperature were observed (data not shown). We then sought to exploit an alternative crystallography-based approach, to verify interaction and to determine binding modes. Ligand soaking was performed for all compounds, however only 10 crystals with **1, 2, 3, 9, 10, 11, 12, 14, 15** and **19** still preserved diffraction quality. Examination of electron density maps revealed in all structures additional density in proximity to Y78 backbone where the carbonyl of bound acetyl-lysine was typically situated (Figure 3 and Supplementary Figure S3). This consistent observation likely confirmed our hypotheses of the use of a central amide group as an acetyl-lysine mimetic for ENL. However, an assessment of the ligand binding modes was only possible for compound **12** and **19**, where a complete trace of electron density enabled an accurate placement of the ligands (Figure 3D). These compounds were from two different groups in our selection yet their binding modes were similar. First, the amide core was observed to flip in comparison to that of acetyl-lysine. While the carbonyl of the flipped amide maintained the β-sheet type hydrogen bonding pattern of acylated lysine to Y78 backbone, the nitrogen group further engaged a direct bond to the S58 side chain that swung slightly backwards. At the front end of the ENL pocket both aromatic benzene and benzimidazole groups of **12** and **19**, respectively, attached to the amide carbonyl at R^1^ were sandwiched between loop4 H56 and loop 6 A79, adopting a nearly planar orientation to the histidine imidazole. The decorations of the R^1^ aromatic groups of both compounds were highly solvent exposed, showing little or no interaction to the protein.

Surprisingly, accommodation of the R^2^ decoration at the rear pocket were highly different between these two ligands. For **12**, the R^2^ benzene ring directly attached on to the amide nitrogen atom tucked in planar in between F28 and F59, and potentially formed three-layer μ-stacking with these phenylalanine residues, an interaction expected to mimic lysine crotonylation. In contrast, a similar interaction was not observed for the chlorobenzene of **19**. This could be due to a constraint geometry of the sp3 carbon spacing atom at R^2^, which in comparison to **12** forced a rotation of nearly 120° to ascend the chlorobenzene group outwards from the rear pocket to the solvent exposed region. The orientation of the ring system of **19** potentially also created steric constraints for Y78, and consequently induced the tyrosine to adopt the ‘out’ conformation. Unlike **12**, this conformation of Y78 resulted in a rather exposed, less rigid pocket. Both co-crystal structures of ENL with **12** and **19** provided valuable structural insights into the binding mode highlighting for example the importance of direct attachments of the aromatic moieties to the R^1^ and R^2^ position of the amide core, which may provide fundamental keys not only for a decrease of flexibility of both ligands and Y78 but for enhancing optimal conditions for strong π-π contacts. In parallel, development of a high-throughput assay and screening identified similar chemotypes, in essence a benzimidazole-amide hit with low micromolar affinity^30^.

To test this chemical scaffold, we used our available chemistry to preliminarily synthesize a small set of R^2^-benzimidazole amide derivatives with modification on R^1^, resulting in compounds **20-24** (Figure 4A). Using ITC, we observed that these derivatives bound ENL with good affinities in low micromolar range with compound **20** showing the best potency demonstrating a submicromolar *K*_d_ of 0.8 μM. We then focused on characterizing of the binding mode of **20**, and successful soaking and structure determination enabled an insight into the binding mode of this compound in ENL. As expected the Kac mimetic amide core retained its flipped binding orientation maintaining the interactions to Y78 and S58 as observed for **12** and **19** (Figure 4B). The R^1^ 3-iodo-4-methylbenzene was situated in the front cavity, adopting a planar conformation to H56 for a μ-stacking contact (distance ^∼^4 Å). The R^2^ benzimidazole as expected located at the rear pocket was sandwiched between and feasibly induced a three-layer π-stacking with F28 and F59. However, apart from these predictable fundamental key interactions compound **20** was observed to engage further hydrogen bonds to the protein, which were not present previously in **12** and **19**. This included a direct contact between the nitrogen of the benzimidazole ring and S76 backbone carbonyl. In addition, the extended piperidine decoration was observed to protrude further along the protein surface at the rear end, where it exerted a conformation that enabled a direct contact between its nitrogen with the E75 carboxylic side chain (distance of ^∼^3.1 Å). We performed ITC to assess thermodynamics of the binding of **12, 19** and **20** to ENL (Figure 4C). Binding of **20** was characterized by a favorable enthalpic binding enthalpy and a *K*_d_of ^∼^807 nM. Binding enthalpy was observed for **12** and **19** indicating interactions in the 20-50 ⍰*K*_d_ region. The presented data demonstrate that the acetyl-lysine binding site of ELN is druggable using a central aromatic scaffold decorated with an amide that acts as acetyl-lysine mimetic moiety.

**Figure 4.**
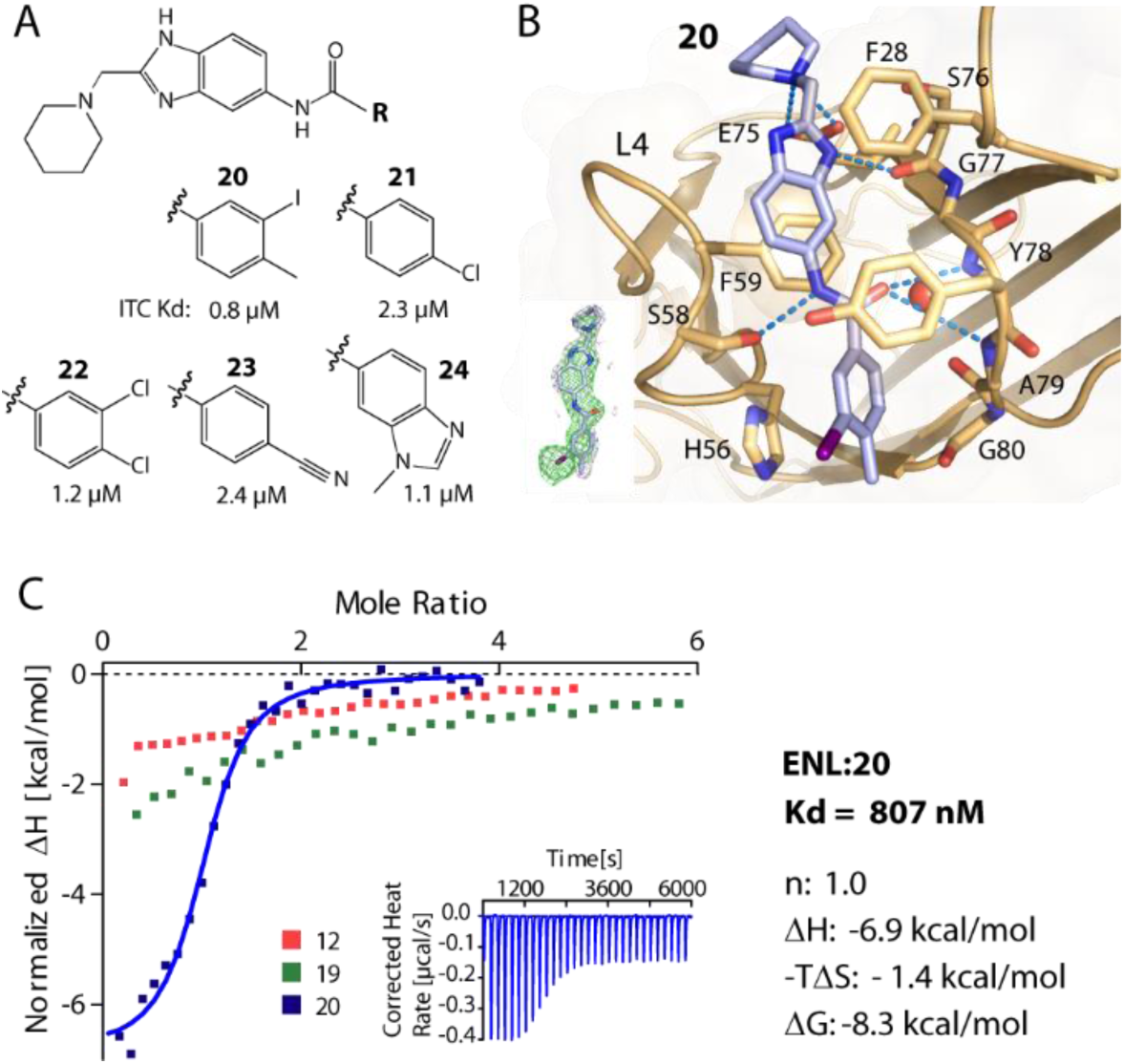
Benzimidazole-amide derivatives and characterization of the interaction of **20** with ENL. Å) Chemical structures of a set of benzimidazole-amide derivatives and their ITC Kd values. B) Crystal structure of the ENL complex with **20** (pdb code, 6hpw). Insets show |*F*_O_| - |*F*_C_| omitted electron density map contoured at 3σ (green) and |2*F*_O_| - |*F*_C_| refined map contoured at 1σ (grey). C) ITC normalized binding heat of the interactions between **12**, **19** and **20** with ENL. Inset shows the isotherm of raw injection heats for **20**.

YEATS domains have a fundamentally different binding site than the well-explored acetyl-lysine readers of the bromodomain family. These differences enable YEATS domains to also recognize larger modifications at the ε-nitrogen of lysines including crotonylation and others acyl-modifications not recognized by bromodomains which typically have a closed water-filled binding pocket. Thus, targeting YEATS domains will require a different design of acyl-lysine mimetic inhibitors. The information provided by this structure-based fragment study will facilitate this design and the development of more potent inhibitors. In our laboratory, we used benzimidazole amides for the development of a potent chemical probe for ELN and AF9^31^, which will facilitate further study of the role of this interesting reader domain in normal physiology and disease.

## Experimental section

Protein purification, crystallization and structure determination - ENL YEATS domain (1-148) was sub-cloned into pNIC-CH, and the recombinant protein incorporating a non-cleavable C-terminal His_6_-tag was expressed *E. coli* Rosetta. Purification was performed using Ni^2+^ affinity and size-exclusion chromatography. Pure protein in 25 mM Tris pH 7.5, 300 mM NaCl, 0.2 mM TCEP was concentrated to 0.45 mM, and was used for crystallization at 20 °C and various reservoir solutions using PEG/PEG smears^32^ as precipitant (Supplementary table S1). Ligands at 5-40 mM concentration were soaked into crystals, which were subsequently cryo-protect with 25% ethylene glycol. Diffraction data collected at BESSY II, SLS, and Diamond Light Source were processed using XDS^33^ or iMOSFLM^34^ and scaled with aimless^35^. Molecular replacement was performed in PHASER^36^ using pdb-id: 5J9s. Model rebuilding alternated with structure refinement was performed in COOT^37^ and REFMAC^38^. The models were verified using molprobity^39^. Data collection and refinement statistics are summarized in Suppl. table S1.

Compounds used in this study were obtained from commercial source (ChemDiv (1-7, 9-10, 13-15, 17-19), Enamine (8 and 16)), except 11, **12, 20-24**, which were synthesized and checked for their purity of >95% using either the stated Waters UHPLC or the Varian ProStar HPLC. Detailed syntheses and characterizations are provided in Supplementary information.

Isothermal titration calorimetry data were measured with a NanoITC instrument (TA instruments) at 25°C. Protein at 300 μM in 25 mM Tris pH 7.5, 500 mM NaCl, 0.5 TCEP, 5% glycerol was titrated into 20-30 μM compounds.

## Author Contributions

AC and SK wrote the paper with contributions of all authors. DH conducted the experiments, MM, DM and JS contributed inhibitors, PB, OF, SK, AC supervised research. All authors have given approval to the final version of the manuscript.

## Notes

Coordinates and structure factors have been deposited: accession codes 6hq0, 6hpw, 6hpx, 6hpy and 6hpz.

## ACKNOWLEDGMENT

The SGC, a registered charity that receives funds from AbbVie, Bayer Pharma AG, Boehringer Ingelheim, Canada Foundation for Innovation, Eshelman Institute for Innovation, Genome Canada, Innovative Medicines Initiative [ULTRA-DD 115766], Wellcome Trust, Janssen, Merck & Co., Novartis Pharma AG, Ontario Ministry of Economic Development and Innovation, Pfizer, São Paulo Research Foundation-FAPESP, Takeda and the Centre of Excellence Macromolecular complexes (CEF). S.K. and A.C. are supported by the Sonderforschungsbereich SFB1177 Autophagy. M.M. is grateful to the EPSRC Centre for Doctoral Training in Synthesis for Biology and Medicine (EP/L015838/1) for a studentship, supported by AstraZeneca, Diamond, Defence Science and Technology Laboratory, Evotec, GlaxoSmithKline, Janssen,
Novartis, Pfizer, Syngenta, Takeda, UCB and Vertex. The authors thank staffs at BESSY II, SLS and Diamond Light Source for their support.

## ABBREVIATIONS

YEATS domain: The Yaf9, ENL, AF9, Taf14, Sas5 (YEATS) domain
ENL: eleven-nineteen-leukemia protein
MLLT1: myeloid/lymphoid or mixed-lineage leukemia translocated to, chromosome 1 protein
AF9: ALL1-fused gene from chromosome 9 protein
MLLT3: myeloid/lymphoid or mixed-lineage leukemia translocated to chromosome 3 protein

## REFERENCES

1. Dhalluin, C.; Carlson, J. E.; Zeng, L.; He, C.; Aggarwal, A. K.; Zhou, M. M., Structure and ligand of a histone acetyltransferase bromodomain. Nature 1999, 399 (6735), 491–496.

2. Filippakopoulos, P.; Picaud, S.; Mangos, M.; Keates, T.; Lambert, J. P.; Barsyte-Lovejoy, D.; Felletar, I.; Volkmer, R.; Muller, S.; Pawson, T.; Gingras, A. C.; Arrowsmith, C. H.; Knapp, S., Histone recognition and large-scale structural analysis of the human bromodomain family. Cell 2012, 149 (1), 214–231.

3. Li, Y.; Wen, H.; Xi, Y.; Tanaka, K.; Wang, H.; Peng, D.; Ren, Y.; Jin, Q.; Dent, S. Y.; Li, W.; Li, H.; Shi, X., AF9 YEATS domain links histone acetylation to DOT1L-mediated H3K79 methylation. Cell 2014, 159 (3), 558–571.

4. Mi, W.; Guan, H.; Lyu, J.; Zhao, D.; Xi, Y.; Jiang, S.; Andrews, F. H.; Wang, X.; Gagea, M.; Wen, H.; Tora, L.; Dent, S. Y. R.; Kutateladze, T. G.; Li, W.; Li, H.; Shi, X., YEATS2 links histone acetylation to tumorigenesis of non-small cell lung cancer. Nat Commun 2017, 8 (1), 1088.

5. Wan, L.; Wen, H.; Li, Y.; Lyu, J.; Xi, Y.; Hoshii, T.; Joseph, J. K.; Wang, X.; Loh, Y. E.; Erb, M. A.; Souza, A. L.; Bradner, J. E.; Shen, L.; Li, W.; Li, H.; Allis, C. D.; Armstrong, S. A.; Shi, X., ENL links histone acetylation to oncogenic gene expression in acute myeloid leukaemia. Nature 2017, 543 (7644), 265–269.

6. Hsu, C. C.; Zhao, D.; Shi, J.; Peng, D.; Guan, H.; Li, Y.; Huang, Y.; Wen, H.; Li, W.; Li, H.; Shi, X., Gas41 links histone acetylation to H2A.Z deposition and maintenance of embryonic stem cell identity. Cell Discov 2018, 4, 28, 3–17.

7. Li, Y.; Sabari, B. R.; Panchenko, T.; Wen, H.; Zhao, D.; Guan, H.; Wan, L.; Huang, H.; Tang, Z.; Zhao, Y.; Roeder, R. G.; Shi, X.; Allis, C. D.; Li, H., Molecular Coupling of Histone Crotonylation and Active Transcription by AF9 YEATS Domain. Mol Cell 2016, 62 (2), 181–193.

8. Andrews, F. H.; Shinsky, S. A.; Shanle, E. K.; Bridgers, J. B.; Gest, A.; Tsun, I. K.; Krajewski, K.; Shi, X.; Strahl, B. D.; Kutateladze, T. G., The Taf14 YEATS domain is a reader of histone crotonylation. Nat Chem Biol 2016, 12 (6), 396–398.

9. Zhao, D.; Guan, H.; Zhao, S.; Mi, W.; Wen, H.; Li, Y.; Zhao, Y.; Allis, C. D.; Shi, X.; Li, H., YEATS2 is a selective histone crotonylation reader. Cell Res 2016, 26 (5), 629–632.

10. Le Masson, I.; Yu, D. Y.; Jensen, K.; Chevalier, A.; Courbeyrette, R.; Boulard, Y.; Smith, M. M.; Mann, C., Yaf9, a novel NuA4 histone acetyltransferase subunit, is required for the cellular response to spindle stress in yeast. Mol Cell Biol 2003, 23 (17), 6086–6102.

11. Wang, A. Y.; Schulze, J. M.; Skordalakes, E.; Gin, J. W.; Berger, J. M.; Rine, J.; Kobor, M. S., Asf1-like structure of the conserved Yaf9 YEATS domain and role in H2A.Z deposition and acetylation. Proc Natl Acad Sci U S A 2009, 106 (51), 21573–21578.

12. Schulze, J. M.; Wang, A. Y.; Kobor, M. S., YEATS domain proteins: a diverse family with many links to chromatin modification and transcription. Biochem Cell Biol 2009, 87 (1), 65–75.

13. Andrews, F. H.; Shanle, E. K.; Strahl, B. D.; Kutateladze, T. G., The essential role of acetyllysine binding by the YEATS domain in transcriptional regulation. Transcription 2016, 7 (1), 14–20.

14. Shanle, E. K.; Andrews, F. H.; Meriesh, H.; McDaniel, S. L.; Dronamraju, R.; DiFiore, J. V.; Jha, D.; Wozniak, G. G.; Bridgers, J. B.; Kerschner, J. L.; Krajewski, K.; Martin, G. M.; Morrison, A. J.; Kutateladze, T. G.; Strahl, B. D., Association of Taf14 with acetylated histone H3 directs gene transcription and the DNA damage response. Genes Dev 2015, 29 (17), 1795–1800.

15. Zhao, D.; Li, Y.; Xiong, X.; Chen, Z.; Li, H., YEATS Domain-A Histone Acylation Reader in Health and Disease. J Mol Biol 2017, 429 (13), 1994–2002.

16. Dutta, A.; Abmayr, S. M.; Workman, J. L., Diverse Activities of Histone Acylations Connect Metabolism to Chromatin Function. Mol Cell 2016, 63 (4), 547–552.

17. Erb, M. A.; Scott, T. G.; Li, B. E.; Xie, H.; Paulk, J.; Seo, H. S.; Souza, A.; Roberts, J. M.; Dastjerdi, S.; Buckley, D. L.; Sanjana, N. E.; Shalem, O.; Nabet, B.; Zeid, R.; Offei-Addo, N. K.; Dhe-Paganon, S.; Zhang, F.; Orkin, S. H.; Winter, G. E.; Bradner, J. E., Transcription control by the ENL YEATS domain in acute leukaemia. Nature 2017, 543 (7644), 270–274.

18. Zeisig, D. T.; Bittner, C. B.; Zeisig, B. B.; Garcia-Cuellar, M. P.; Hess, J. L.; Slany, R. K., The eleven-nineteen-leukemia protein ENL connects nuclear MLL fusion partners with chromatin. Oncogene 2005, 24 (35), 5525–5532.

19. Heisel, S.; Habel, N. C.; Schuetz, N.; Ruggieri, A.; Meese, E., The YEATS family member GAS41 interacts with the general transcription factor TFIIF. BMC Mol Biol 2010, 11, 53.

20. Filippakopoulos, P.; Knapp, S., Targeting bromodomains: epigenetic readers of lysine acetylation. Nat Rev Drug Discov 2014, 13 (5), 337–356.

21. Hewings, D. S.; Rooney, T. P.; Jennings, L. E.; Hay, D. A.; Schofield, C. J.; Brennan, P. E.; Knapp, S.; Conway, S. J., Progress in the development and application of small molecule inhibitors of bromodomain-acetyl-lysine interactions. J Med Chem 2012, 55 (22), 9393–9413.

22. Filippakopoulos, P.; Qi, J.; Picaud, S.; Shen, Y.; Smith, W. B.; Fedorov, O.; Morse, E. M.; Keates, T.; Hickman, T. T.; Felletar, I.; Philpott, M.; Munro, S.; McKeown, M. R.; Wang, Y.; Christie, A. L.; West, N.; Cameron, M. J.; Schwartz, B.; Heightman, T. D.; La Thangue, N.; French, C. A.; Wiest, O.; Kung, A. L.; Knapp, S.; Bradner, J. E., Selective inhibition of BET bromodomains. Nature 2010, 468 (7327), 1067–1073.

23. Vidler, L. R.; Brown, N.; Knapp, S.; Hoelder, S., Druggability analysis and structural classification of bromodomain acetyl-lysine binding sites. J Med Chem 2012, 55 (17), 7346–7359.

24. Ferguson, F. M.; Fedorov, O.; Chaikuad, A.; Philpott, M.; Muniz, J. R.; Felletar, I.; von Delft, F.; Heightman, T.; Knapp, S.; Abell, C.; Ciulli, A., Targeting low-druggability bromodomains: fragment based screening and inhibitor design against the BAZ2B bromodomain. J Med Chem 2013, 56 (24), 10183–10187.

25. Chaikuad, A.; Lang, S.; Brennan, P. E.; Temperini, C.; Fedorov, O.; Hollander, J.; Nachane, R.; Abell, C.; Muller, S.; Siegal, G.; Knapp, S., Structure-Based Identification of Inhibitory Fragments Targeting the p300/CBP-Associated Factor Bromodomain. J Med Chem 2016, 59 (4), 1648–1653.

26. Chaikuad, A.; Petros, A. M.; Fedorov, O.; Xu, J.; Knapp, S., Structure-based approaches towards identification of fragments for the low-druggability ATAD2 bromodomain. Medchemcomm 2014, 5 (12), 1843–1848.

27. Filippakopoulos, P.; Knapp, S., The bromodomain interaction module. FEBS Lett 2012, 586 (17), 2692–2704.

28. Aldeghi, M.; Ross, G. A.; Bodkin, M. J.; Essex, J. W.; Knapp, S.; Biggin, P. C., Large-scale analysis of water stability in bromodomain binding pockets with grand canonical Monte Carlo. Commun Chem 2018, 1–24.

29. Aldeghi, M.; Bodkin, M. J.; Knapp, S.; Biggin, P. C., Statistical Analysis on the Performance of Molecular Mechanics Poisson-Boltzmann Surface Area versus Absolute Binding Free Energy Calculations: Bromodomains as a Case Study. J Chem Inf Model 2017, 57 (9), 2203–2221.

30. Christott, T.; Bennett, J.; Coxon, C.; Monteiro, O.; Giroud, C.; Beke, V.; Felce, S. L.; Gamble, V.; Gileadi, C.; Poda, G.; Al-Awar, R.; Farnie, G.; Fedorov, O., Discovery of a Selective Inhibitor for the YEATS Domains of ENL/AF9. SLAS Discov 2018, 2472555218809904.

31. Moustakim, M.; Christott, T.; Monteiro, O. P.; Bennett, J.; Giroud, C.; Ward, J.; Rogers, C. M.; Smith, P.; Panagakou, I.; Saez, L. D.; Felce, S. L.; Gamble, V.; Gileadi, C.; Halidi, N.; Heidenreich, D.; Chaikuad, A.; Knapp, S.; Huber, K. V. M.; Farnie, G.; Heer, J.; Manevski, N.; Poda, G.; Al-Awar, R.; Dixon, D. J.; Fedorov, O.; Brennan, P., Discovery of an MLLT1/3 YEATS Domain Chemical Probe. Angew Chem Int Ed Engl 2018. [Epub ahead of print]

32. Chaikuad, A.; Knapp, S.; von Delft, F., Defined PEG smears as an alternative approach to enhance the search for crystallization conditions and crystal-quality improvement in reduced screens. Acta Crystallogr D Biol Crystallogr 2015, 71 (Pt 8), 1627–1639.

33. Kabsch, W., Xds. Acta Crystallogr D Biol Crystallogr 2010, 66 (Pt 2), 125–132.

34. Powell, H. R.; Battye, T. G. G.; Kontogiannis, L.; Johnson, O.; Leslie, A. G. W., Integrating macromolecular X-ray diffraction data with the graphical user interface iMosflm. Nat Protoc 2017, 12 (7), 1310–1325.

35. Potterton, L.; Agirre, J.; Ballard, C.; Cowtan, K.; Dodson, E.; Evans, P. R.; Jenkins, H. T.; Keegan, R.; Krissinel, E.; Stevenson, K.; Lebedev, A.; McNicholas, S. J.; Nicholls, R. A.; Noble, M.; Pannu, N. S.; Roth, C.; Sheldrick, G.; Skubak, P.; Turkenburg, J.; Uski, V.; von Delft, F.; Waterman, D.; Wilson, K.; Winn, M.; Wojdyr, M., CCP4i2: the new graphical user interface to the CCP4 program suite. Acta Crystallogr D Struct Biol 2018, 74 (Pt 2), 68–84.

36. McCoy, A. J., Acknowledging Errors: Advanced Molecular Replacement with Phaser. Methods Mol Biol 2017, 1607, 421–453.

37. Emsley, P., Tools for ligand validation in Coot. Acta Crystallogr D Struct Biol 2017, 73 (Pt 3), 203–210.

38. Vagin, A. A.; Steiner, R. A.; Lebedev, A. A.; Potterton, L.; McNicholas, S.; Long, F.; Murshudov, G. N., REFMAC5 dictionary: organization of prior chemical knowledge and guidelines for its use. Acta Crystallogr D Biol Crystallogr 2004, 60 (Pt 12 Pt 1), 2184–2195.

39. Williams, C. J.; Headd, J. J.; Moriarty, N. W.; Prisant, M. G.; Videau, L. L.; Deis, L. N.; Verma, V.; Keedy, D. A.; Hintze, B. J.; Chen, V. B.; Jain, S.; Lewis, S. M.; Arendall, W. B., 3rd; Snoeyink, J.; Adams, P. D.; Lovell, S. C.; Richardson, J. S.; Richardson, D. C., MolProbity: More and better reference data for improved all-atom structure validation. Protein Sci 2018, 27 (1), 293–315.

